# Python nidoviruses, more than respiratory pathogens

**DOI:** 10.1101/2020.04.10.036640

**Authors:** Eva Dervas, Jussi Hepojoki, Teemu Smura, Barbara Prähauser, Katharina Windbichler, Sandra Blümich, Antonio Ramis, Udo Hetzel, Anja Kipar

## Abstract

In recent years nidoviruses have emerged as an important respiratory pathogen of reptiles, affecting especially captive python populations. In pythons, nidovirus infection induces an inflammation of the upper respiratory and alimentary tract which can develop into a severe and often fatal proliferative pneumonia. We observed pyogranulomatous and fibrinonecrotic lesions in organ systems other than the respiratory tract during full post mortem examinations on 30 nidovirus RT-PCR positive pythons of varying species originating from Switzerland and Spain. The observations prompted us to study whether the atypical tissue tropism associates with previously unknown nidoviruses or changes in the nidovirus genome. RT-PCR and inoculation of *Morelia viridis* cell cultures served to recruit the cases and to obtain virus isolates. Immunohistochemistry and immunofluorescence staining against nidovirus nucleoprotein demonstrated that the virus not only infects a broad spectrum of epithelial (respiratory and alimentary epithelium, hepatocytes, renal tubules, pancreatic ducts etc.), but also intravascular monocytes, intralesional macrophages and endothelial cells. By next-generation sequencing we obtained full length genome for a novel nidovirus species circulating in Switzerland. Analysis of viral genomes recovered from pythons showing nidovirus infection-associated respiratory or systemic disease did not explain the observed phenotypes. The results indicate that python nidoviruses have a broad cell and tissue tropism, further suggesting that the course of infection could vary and involve lesions in a broad spectrum of tissues and organ systems as a consequence of monocyte-mediated systemic spread of the virus.

**IMPORTANCE:** During the last years, python nidoviruses have become a primary cause of fatal disease in pythons. Nidoviruses represent a threat to captive snake collections, as they spread rapidly and can be associated with high morbidity and mortality. Our study indicates that, different from previously evidence, the viruses do not only affect the respiratory tract, but can spread in the entire body with blood monocytes, have a broad spectrum of target cells, and can induce a variety of lesions. Nidovirales is an order of animal and human viruses that compromise important zoonotic pathogens such as MERS-CoV and SARS-CoV, as well as the recently emerged SARS-CoV-2. Python nidoviruses belong to the same subfamily as the mentioned human viruses and show similar characteristics (rapid spread, respiratory and gastrointestinal tropism, etc.). The present study confirms the relevance of natural animal diseases to better understand the complexity of viruses of the order nidovirales.

## INTRODUCTION

In the past, toroviruses, a subfamily of the order *Nidovirales*, were mainly known to cause enteric disease in mammals (1–4). Recent studies linked nidovirus infections to respiratory disease in cattle, pythons, and lizards, thus establishing nidoviruses as respiratory pathogens (5–12). In pythons, nidoviruses were found to be associated with chronic proliferative pneumonia (5, 6, 10, 11), and the association was confirmed by experimental infection studies (7). Severe cases exhibit typical pathological changes both after experimental and natural infection; these include stomatitis, rhinitis, tracheitis, and pneumonia with significant mucus accumulation. Taxonomically, python nidoviruses have recently been reclassified into the family *Tobaniviridae*, subfamily *Serpentovirinae*, genus *Pregotovirus* (International Committee on Taxonomy of Viruses (ICTV), https://talk.ictvonline.org (July 2018)).

74 A recent study identified yet another new novel nidovirus (genus *Barnivirus*) in Bellinger River snapping turtles *(Myuchelys georgesi).* The infection was associated with necrotizing cystitis, nephritis, adenitis and vasculitis, and the presence of viral RNA in many tissues, indicating systemic spread of the virus (12). The target cell spectrum of python nidoviruses includes the epithelium of the respiratory tract and lungs (6), and in some cases also the oral cavity and the cranial esophagus (5–7, 11), the mucosa of which exhibits ciliated epithelium in snakes (13). Thus far little is known about the intra- and interspecies transmission of python nidoviruses. However, recent studies demonstrated nidovirus RNA in oral and cloacal swabs and the intestinal content of diseased and healthy snakes, suggesting that both airborne and fecal-oral transmission may occur (7, 14, 9–11).

In an earlier report, we studied the pathogenesis of nidovirus-associated pneumonia in green tree pythons (6). Similarly to other investigators who described nidovirus pneumonia in pythons (5, 8, 10, 11), we occasionally detected the virus also in other organs, with and without evidence of pathological changes (6), suggesting that the cell tropism of python nidoviruses goes beyond the respiratory epithelium. These findings and the fact that nidovirus infections affect several python species led us to the hypothesis that python nidoviruses are not restricted to a certain species and have a broad disease potential. Therefore, we undertook a larger study, making use of diagnostic cases with natural nidovirus infection. A total of 30 nidovirus infected snakes, selected based on demonstration of nidovirus nucleoprotein (NP) by immunohistology in tissues with lesions. Six python species from nine breeding colonies and collections were included, namenly the green tree python (*Morelia viridis)*, woma python (*Aspidites ramsayi)*, carpet python (*Morelia spilota*), Angolan python (*Python anchietae*), ball python (*Python regius*), Indian python (*Python molurus*), and the black-headed python (*Aspidites melanocephalus*). Immunohistology also helped to identify the nidovirus target cells (6), and next-generation sequencing (NGS) served to obtain complete or near complete genomes of the causative viruses. The results support the hypothesis that nidovirus infections can cause variable disease in pythons, with lesions in a broad spectrum of tissues and organs, and monocyte-mediated systemic spread of the virus.

## MATERIALS AND METHODS

### Animals

The study included 30 pythons of six different species (green tree python, *Morelia viridis*; woma python, *Aspidites ramsayi*; carpet python, *Morelia spilota*; Angolan python, *Python anchietae;* ball python, *Python regius*; Indian python, *Python molurus*; Black-headed python, *Aspidites melanocephalus*) with nidovirus-associated disease, confirmed by immunohistology for nidovirus nucleoprotein (NP) and reverse transcription-polymerase chain reaction (RT-PCR) (6) (Table 1). The animals had been submitted for a diagnostic post mortem examination upon the owners’ request, either at the Institute of Veterinary Pathology, Vetsuisse Faculty, University of Zurich, or at the Pathology Unit, Universitat Autònoma de Barcelona, Spain, between 2012 and 2018. Most animals had died or been euthanized prior to submission; four individuals (CH-A4, CH-B1, CH-B2, E-B2) were submitted by the owner for euthanasia and immediate diagnostic examination. Euthanasia followed an ASPA (Animals Scientific Procedures Act 1986) schedule 1 (appropriate methods of humane killing, http://www.legislation.gov.uk/ukpga/1986/14/schedule/1) procedure. A separate research permit was not required for the diagnosis-motivated necropsies and subsequent sample collection.

**Table 1.** Animals, clinical signs, lesions and viral target cells. All animals were from breeding collections in Switzerland (CH) or Spain (E). Breeders CH A-C and CH-F were working closely together, trading/exchanging snakes.

| Animal | Species | Age | Sex | Clinical history | Lesions (affected tissues) | Viral target cells | Virus detection |
| --- | --- | --- | --- | --- | --- | --- | --- |
| CH-A1<sup>1</sup> | *Morelia viridis* | 8 y | M | RD, mucus in OC | NPD (NC,T,L), FNI (Sp, Es) | EP (NC, OC, T, L, R, Es) | PCR (L) |
| CH-A2 | *Morelia viridis* | 5 y | M | RD | NPD (T,L) | EP (T, L) | PCR (L) |
| CH-A3 | *Morelia viridis* | 2 y | M | RD | NPD (NC, OC, L) | EP (NC, OC, L) | PCR (L) |
| CH-A4 | *Aspidites ramsayi* | 3 mo | nk | RD, mucus in OC | NPD (NC, OC, L) | EP (NC, OC, L) | PCR (L), NGS (SNT) |
| CH-A5 | *Morelia spilota* | 4 y | F | No history | NPD (NC, OC, T, L) | EP (NC, OC, T, L) | PCR (L), NGS (SNT) |
| CH-A6 | *Morelia spilota* | 1 y | M | RD | NPD (T, L) | EP (T, L) | PCR (Ch, Cl) |
| CH-A7 | *Morelia viridis* | adult | F | Sudden death | NPD (NC, OC, T), FNI (Th, Sp), V (He, Sp, SI), SGD (H, Es, SI) | EP (NC, OC, T, L, RT, H, PD, E, Es, SI), macrophages | PCR (L), NGS (SNT, liver) |
| CH-A8 | *Morelia viridis* | 1.5 y | M | Sudden death | NPD (NC, OC, T), SGD (Es, SI, LI, H, Sp) | EP (NC, OC, T, L, RT, H, PD), VE, macrophages, monocytes | NGS (SNT, liver) |
| CH-A9 | *Python regius* | adult | nk | No history | NPD (NC, OC, T), FNI (Es) | EP (NC, OC, T, L, Es, RT) | PCR (L, Ch, Cl) |
| CH-B1 | *Morelia spilota* | 1 y | M | RD | NPD (NC, OC), FNI (K) | EP (NC, OC, T, L, PD, RT) | PCR (L, Cl), NGS (SNT) |
| CH-B2 | *Morelia spilota* | 1 y | M | RD | NPD (NC) | EP (NC) | PCR (L) |
| CH-B3 | *Morelia viridis* | nk | M | RD | NPD (T, L), FNI (K, P) | EP (L, PD, RT) | PCR (L, Ch, Cl), NGS (SNT) |
| CH-B4 | *Morelia spilota* | 1 y | M | RD | NPD (NC, OC, T, L) | EP (NC, OC, T, L) | PCR (L, Ch) |
| CH-B5<sup>1</sup> | *Python anchietae* | 8 y | M | RD | NPD (T, L) | EP (NC, OC, T, L) | NGS (SNT) |
| CH-B6 | *Morelia viridis* | 1.5 y | M | Sudden death | NPD (NC, OC, T), FNI (Es, SI, LI, Sp), SGD (Es, SI, LI, Th, K, He, H), V (Th, He) | EP (NC, OC, T, L, RT, Es, S, PD); ependyma | PCR (L, K), NGS (liver) |
| CH-C1 | *Morelia viridis* | adult | F | RD | NPD (OC, T, L), SGD (R) | EP (NC, OC, T, L, RT) | PCR (Ch, Cl) |
| CH-C2<sup>1</sup> | *Morelia spilota* | 6 y | F | RD | NPD (NC, OC, T, L) | EP (NC, OC, T, L) | PCR (L) |
| CH-C3 | *Morelia viridis* | juv | F | Sudden death | SGD (L, H, K, SI, LI), FNI (Es, M), V (SI, LI) | EP (L, RT, H, Es, SI, LI, M), VE, macrophages, monocytes | NGS (SNT, liver) |
| CH-C4 | *Aspidites melanocephalus* | adult | M | Anorexia | NPD (T, L), FNI (Es) | EP (T, L, Es) | PCR (L, Ch, Cl) |
| CH-D1 | *Python regius* | nk | M | RD | NPD (NC, OC, T, L), FNI (Es) | EP (T, L, Es) | PCR (L, Ch, Cl) |
| CH-D2 | *Python regius* | nk | M | RD | NPD (NC, OC, T, L), FNI (Es) | EP (T, L, Es) | PCR (L, Ch, Cl), NGS (SNT) |
| CH-E2 | *Python regius* | nk | F | RD | NPD (T, L), FNI (Es) | EP (T, L, Es) | PCR (L, Ch, Cl), NGS (SNT) |
| CH-F1 | *Morelia viridis* | 10 y | F | No history | NPD (NC, OC, T, L), FNI (M, SI, LI), V (Ov), SGD (H, Ov) | EP (NC, OC, T, L, H, RT, SI, LI, PD, Ov) VE, ependyma, macrophages, monocytes | PCR (L), NGS (SNT) |
| CH-F2 | *Morelia viridis* | 6 y | F | Anorexia | NPD (NC, OC, T, L) | EP (NC, OC, T, L, Es, PD) | PCR (L), NGS (SNT) |
| E-A1 | *Python regius* | nk | nk | No history | NPD (L) | EP (L) |  |
| E-A2 | *Python anchietae* | nk | nk | Sudden death | NPD (T, L) | EP (T, L) | PCR (L) |
| E-B1 | *Python regius* | juv | M | RD, mucus in OC, anorexia | NPD (T) | EP (T, L, PD) | PCR (L) |
| E-B2<sup>2</sup> | *Python regius* | 1 y | M | RD, mucus in OC | NPD (NC, OC, T, L) | EP (T, L) | PCR (L), NGS (SNT) |
| E-B3 | *Python regius* | juv | M | RD, mucus in OC | NPD (L) | EP (T, L) | PCR (L), NGS (SNT) |
| E-C1 | *Morelia viridis* | 4-5 y | M | No history | NPD (NC, OC, T, L) | NE | PCR (L), NGS (SNT) |

Viral target cells – viral antigen expression in cells (detected by immunohistology) with or without evidence of cytopathic effect.
Virus detection – In all animals, nidovirus-associated disease was confirmed by immunohistology, confirming the presence of virus within the lesions. In addition, RT-PCR for nidovirus (PCR) was performed on tissue samples and/or cloacal (Cl) and choanal (Ch) swabs (Dervas et al., 2017), or next-generation sequencing (NGS) was performed on culture supernatants from *Morelia viridis* cell cultures (Dervas et al., 2017) incubated with lung homogenates (SNT), or from liver tissue (liver).
Anatomical structures and cell types:
OC - oral cavity; NC - nasal cavity; T – trachea; L - lung; He – heart; Es – esophagus; S – stomach; SI - small intestine; LI - Large intestine; K – kidney; Th – thymus; Sp – spleen; Ov – oviduct; M – mesothelium; PD - pancreatic ducts; RT - renal tubules
EP - epithelial cells; PD - pancreatic duct epithels VE - vascular endothelial cells; Hep – hepatocytes
Lesions:
NPD - nidovirus-associated proliferative disease (with variable degree of epithelial hyperplasia and inflammation)
FNI: Fibrinonecrotizing inflammation
SGD: Systemic granulomatous disease

Most animals had presented clinically with respiratory signs, ranging from mild mucus secretion from the oral and nasal cavity to severe acute, recurrent or chronic dyspnea. In some cases, the breeders reported repeated treatment attempts, occasionally for more than one year, prior to the animals’ death, which included frequent nebulising with an antiseptic solution (F10 Antiseptic Solution-Benzalkonium Chloride and Polyhexamethylene Biguanide, Health and Hygiene (Pty) Ltd) and/or antibiotic treatment with commercial broad-spectrum antibiotics such as Enrofloxacin. In other cases, the animals had died suddenly without prior clinical signs (Table 1). Affected snakes had a broad age range (3 months to 10 years; average: 3.65 y). Nineteen were male and ten were female, the sex of one young individual was unknown (undifferentiated genital organs). The animals originated from nine collections/breeding colonies of varying size (from a single snake up to a collection of 50 breeding animals) in Switzerland and Spain (Table 1).

### Sample collection and screening for infectious agents

All animals were subjected to a full post mortem examination and samples from brain, lung, liver and kidney were collected and stored at −80 °C for further analysis. In addition, samples from all major organs and tissues (brain, respiratory tract, liver, kidney, spleen, gastrointestinal tract, reproductive tract, pancreas) were collected and fixed in 4% buffered formalin for histological and immunohistological examinations.

The lungs of some snakes were submitted to routine bacteriological examination, and from one animal, a virological examination for common respiratory snake viruses was undertaken (Table 1).

### Histology, immunohistology and immunofluorescence

Formalin-fixed tissue samples were trimmed and routinely paraffin wax embedded. Consecutive sections (4-5 μm) were prepared and stained with hematoxylin and eosin (HE) or subjected to immunohistological and immunofluorescence staining.

Sections from all histologically examined organs were subjected to immunohistological staining for nidovirus NP, using a custom made rabbit polyclonal antibody (anti-MVNV NP) and a previously described protocol (6). A formalin-fixed, paraffin embedded cell pellet prepared from nidovirus-infected cell cultures served as positive control. Consecutive sections incubated with the pre-immune serum instead of the specific primary antibody served as negative controls. Sections from one individual (CH-A7) underwent immunofluorescence staining. After deparaffinization, they were incubated with anti-MVNV NP antibody at a 1:500 dilution (in phosphate-buffered saline, PBS) overnight at 4 °C; washed five times with PBS, incubated for 30 min at room temperature (RT) with 1:500-diluted (in PBS) Alexa Fluor 594-labeled goat anti-rabbit immunoglobulin secondary antibody (Invitrogen) and then washed four times with PBS. Afterwards, the sections were incubated with an anti-Iba-1 antibody (ab5076, Biotec) at a 1:400 dilution (in PBS) overnight at 4 °C, washed five times with PBS, and incubated for 30 min at RT with Alexa Fluor 488-labeled donkey anti-goat immunoglobulin secondary antibody (Invitrogen; 1:500 in PBS), followed by a 15 min incubation with DAPI (4′, 6-diamidino-2-phenylindole, Novus Biologicals; 1:10,000 in PBS). Sections were washed twice with distilled water, air dried, and a coverslip placed with FluoreGuard mounting medium (Biosystems, Switzerland). Images were taken at a 400 x magnification with a Nikon Eclipse Ti-U inverted microscope with NIS Advanced Research software.

### Virus isolation

Cultured brain and liver cells of *M. viridis*, at passage 8-15, were used for inoculations with tissue samples from selected animals (Table 1) as described (6). At 4-5 days post inoculation, the cell culture supernatants were collected for RT-PCR and next generation sequencing (NGS) library preparation.

### RT-PCR and NGS

RNA was extracted from lung and/or liver tissue samples of 24 animals, from cotton dry swabs used to sample the choanal and cloacal mucosa of 10 animals, from a lung lavage sample of an adult female ball python (*Python regius*) from a further Swiss breeder unrelated to any of the others, and from tissue culture supernatants (Table 1) as described (6). Library preparation, data analysis, and genome assembly was done as described (6).

To study the frequencies of single nucleotide polymorphisms (SNPs) within samples, the NGS reads were quality filtered using Trimmomatic and the reads with Q-score over 30 were assembled against the consensus sequence of a given sample using the BWA-MEM algorithm (15) followed by removal of potential PCR duplicates using SAMTools version 1.8 (16). The frequency of single nucleotide variants in each sequence position was called using LoFreq version 2 (17) and the genetic diversity between viral sequences derived from tissue and cell culture were compared using SNPGenie software (18).

### Phylogenetic analysis

The sequences of the representatives of the genus *Tobaniviridae* were downloaded from the GenBank. The complete genomes of python-associated virus strains were aligned using ClustalW algorithm implemented in the MEGA7 program (19). In addition, the amino acid sequences of ORF1b (RdRp) were aligned with MAFFT version 7.407 using E-INS-i parameters (20). The phylogenetic trees were constructed using the Bayesian Markov chain Monte Carlo (MCMC) method, implemented in Mr. Bayes version 3.2 (21) with two independent runs and four chains per run, the GTR-G-I model of substitution for nucleotides and the WAG model of substitution for amino acids. The analyses were run for 5 million states and sampled every 5,000 steps.

### Recombination analysis

Recombination events were sought from an alignment containing snake-associated nidoviruses using pairwise homoplasy index (PHI) test (22) implemented in SplitsTree version 4.15.1 (23) and tree order scan method implemented in SSE 1.3 software (24), followed by identification of potentially recombinant sequences and estimation of recombination break points using RDP (25), bootscan (26), maxchi (27), chimaera (28), 3seq (29), geneconv (30) and siscan (31) methods implemented in RDP4 software (32). The potential recombination events detected by at least five out of seven methods were further evaluated by constructing phylogenetic trees using neighbor joining method and maximum composite likelihood substitution model implemented in MEGA7 software (19).

## RESULTS

### Gross presentation of nidovirus-associated disease in pythons

Full post mortem examinations were performed on all cases. Most snakes that had died spontaneously exhibited the typical, previously described respiratory changes, represented by a varying amount of mucoid material in the airways and particularly in the lungs (Fig. 1A). In some animals, the oral cavity, the trachea, the lung and the air sacs were filled with mucoid material and the lung parenchyma appeared thickened (nidovirus-associated proliferative disease; Fig.1A); also, the mucoid material was occasionally mixed with purulent exudate. Two euthanized carpet pythons with clinically reported respiratory distress (CH-B1 and -B2, Table 1) did not exhibit any gross pulmonary changes, but showed mild mucus accumulation in the oral cavity. In others, granulomatous and/or fibrinonecrotic lesions were observed in the oral, esophageal and intestinal mucosa (Figs. 1A, B and 2A, B), in liver (Fig. 3A), spleen, or kidney, the coelomic cavity and the oviduct. There was no evidence of a specific lesion pattern in the different affected python species, the findings for each animal are summarized in Table 1.

**Figure 1.**
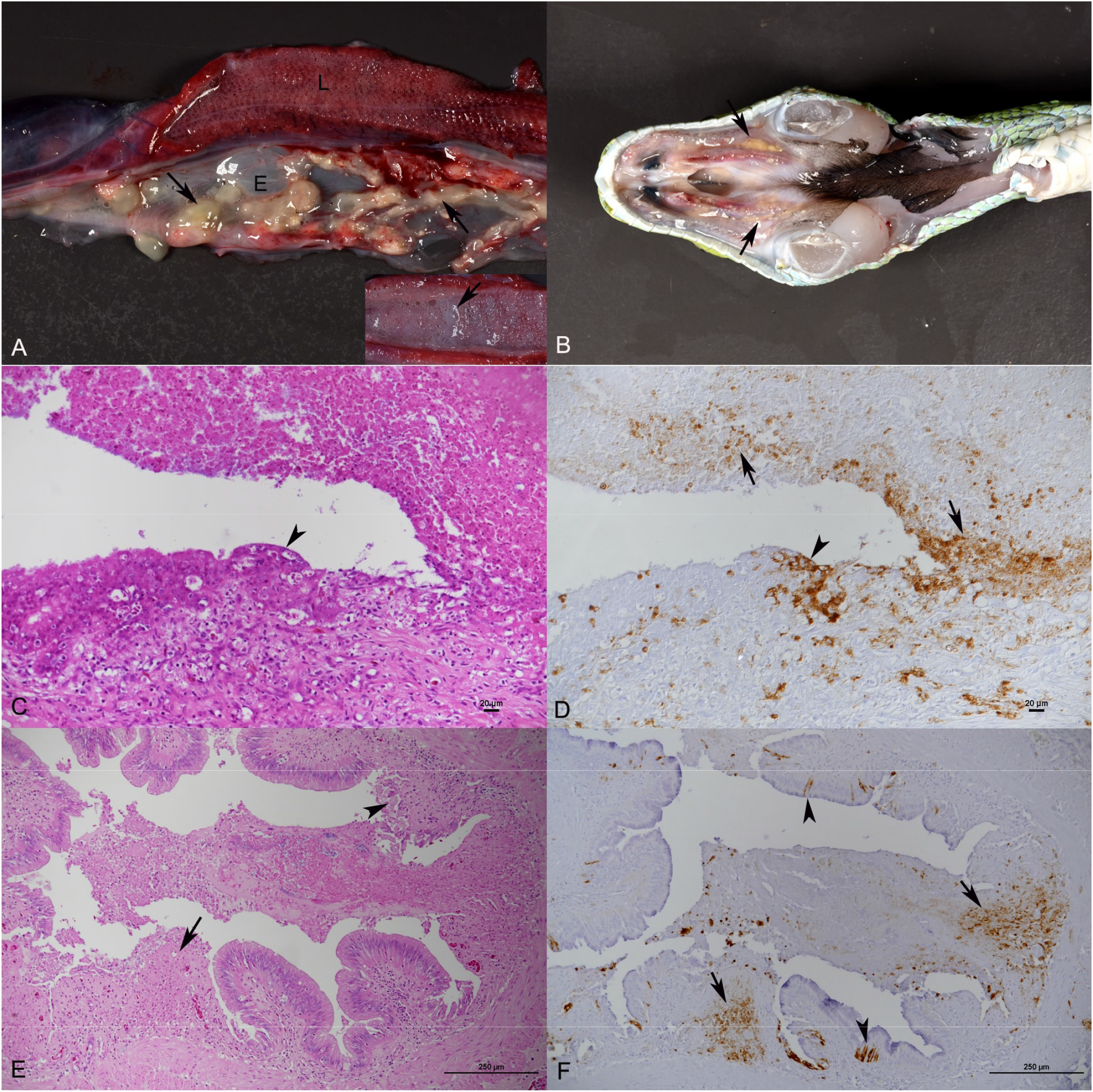
Python nidovirus disease. Lesions in the oral cavity and esophagus. **A, B**. Snake CH-A8 (Green tree python; *Morelia viridis*). **A.** Lung (L) and esophagus (E). The lung appears voluminous and moist. The esophagus exhibits a severe multifocal fibrinonecrotic inflammation (arrow). Inset: lung after longitudinal section, with accumulation of abundant mucoid fluid in the lumen. **B.** Oral cavity, ventral aspect. Focal fibrinonecrotic stomatitis (arrows). **C, D**. Snake CH-A4 (Woma python; *Aspidites ramsayi*). Nasal mucosa. Severe fibrinonecrotic rhinitis (C). There is extensive nidovirus nucleoprotein (NP) expression (D) both cell-free in areas of necrosis (arrows) and within epithelial cells (arrowheads). Bars = 20 μm. **E, F**. Snake CH-A7 (Green tree python; *Morelia viridis*). Esophagus. Severe multifocal fibrinonecrotic esophagitis (E). Nidovirus NP expression (E) is seen both cell-free in areas of necrosis (arrows) and within infected epithelial cells (arrowheads). Bars = 250 μm. C, E: HE stain. D, F: Immunohistology, hemalaun counterstain.

**Figure 2.**
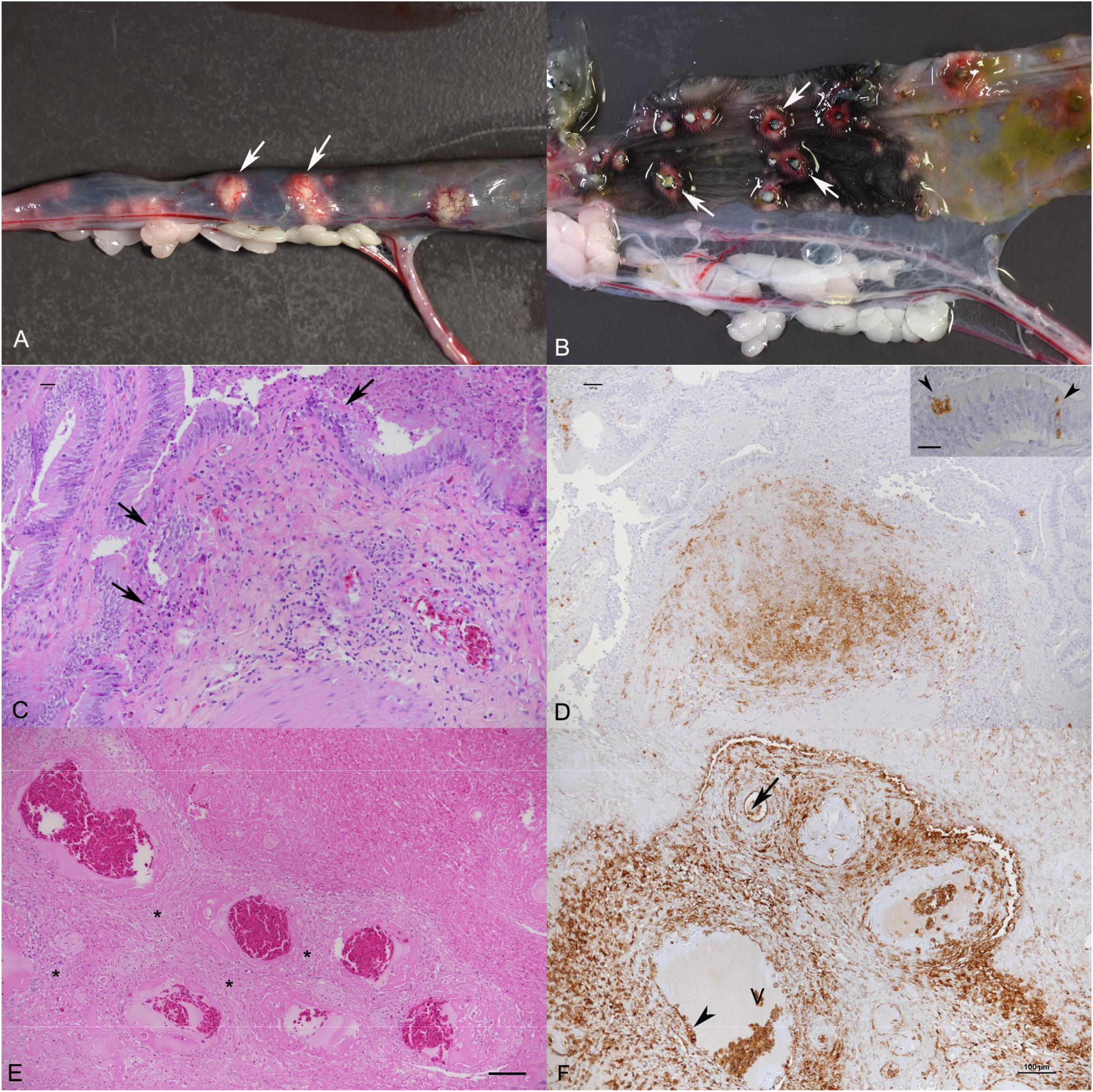
Python nidovirus disease. Intestinal lesions. **A, B**. Snake CH-B6 (Green tree python; *Morelia viridis*). Severe multifocal fibrinonecrotic enteritis with ulcerations (arrows). **C, D**. Snake CH-C3 (Green tree python; *M. viridis*). Small intestine. Severe focal pyogranulomatous perivascular infiltrates and superficial fibrinonecrotic inflammation (C; arrows). Nidovirus NP is abundantly expressed within pyogranulomatous infiltrates (D). Inset: Intact infected enterocytes adjacent to the ulceration (arrows). **E, F**. Snake CH-A7 (Green tree python; *M.*). Large intestine, submucosa. Severe pyogranulomatous perivascular infiltrates (E) with extensive nidovirus NP expression (F) within inflammatory infiltrates and within intravascular mononuclear cells (arrow) and vascular endothelial cells (arrowhead). V: vein. Bars = 100 μm. C, E: HE stain, D, F: Immunohistology, hemalaun counterstain. C: Bar = 20 μm. D: Bar = 50 μm, Inset: Bar = 20 μm.

**Figure 3.**
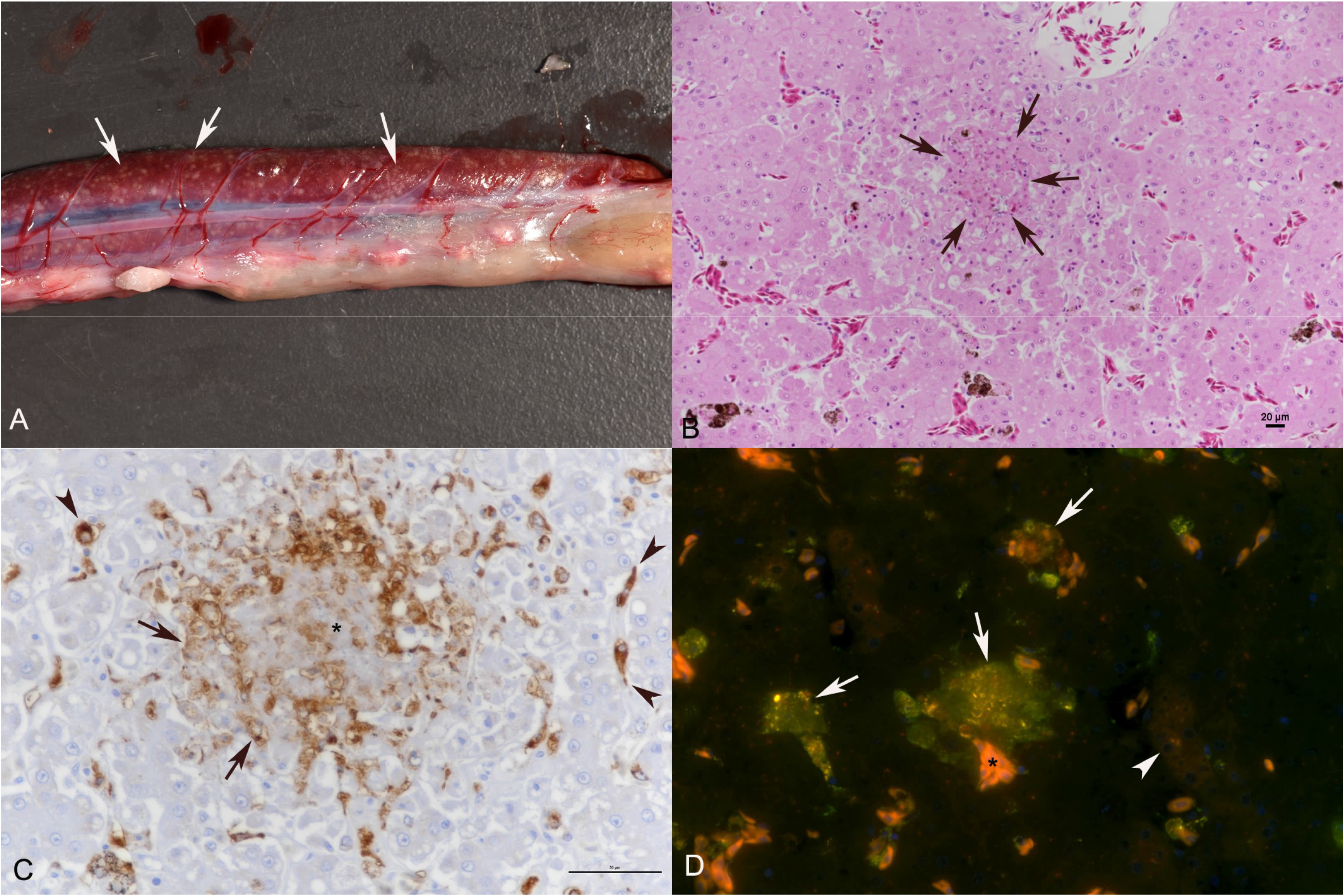
Python nidovirus disease. Involvement of the liver. **A**. Snake CH-B6 (Green tree python; *Morelia viridis*). Severe multifocal pyogranulomatous hepatitis (arrows). **B-D**. Snake CH-C3 (Green tree python; *M. viridis*). **B**. Focal granuloma (arrows). HE stain. Bar = 20 μm. **C**. Closer view of granuloma, with central area of necrosis (asterisk), surrounded by numerous macrophages that exhibit strong nidovirus NP expression (arrows). Viral antigen expression is also seen in Kupffer cells (arrowheads). Immunohistology, hemalaun counterstain. Bar = 50 μm. **D**. Double immunofluorescence of the granuloma confirms nidovirus NP expression (red) within macrophages (Iba-1+; green; arrows) and a Kupffer cell (asterisk). There is also a hepatocyte with weak nidovirus NP expression (arrowhead).

### Python nidoviruses have a broad target cell and lesion spectrum

From all animals, the major organs and tissues as well as gross lesions were examined by histology and immunohistology for nidovirus NP. The examination a revealed broad target cell spectrum and nidovirus-associated cytopathic effect.

#### Oral cavity and respiratory tract

The respiratory tract was found to be affected in all animals (100%), with a variable distribution of viral infection and lesions. In 10 animals (33%), the nasal and oral cavities were affected and exhibited multifocal extensive epithelial necrosis with diffuse subepithelial infiltration of the adjacent, partly hyperplastic respiratory and squamous epithelium by numerous heterophils, lymphocytes, plasma cells and macrophages. Lesions were occasionally covered with fibrin, degenerated epithelial cells and heterophils (fibrinonecrotic rhinitis and stomatitis; Fig. 1C). Nidovirus NP was abundantly expressed in numerous unaltered and degenerated oral and nasal epithelial cells (Fig. 1D). In two euthanized carpet pythons (CH-B1 and -B2, Table 1), these were the only pathological changes, suggesting that they represented early (initial) lesions. The latter is further supported by the fact that in one of the two individuals (CH-B1) nidovirus NP was also detected in a few individual respiratory epithelial cells in the trachea, some trabecular pseudostratified lung epithelial cells, and occasional type I pneumocytes lining the faveolar space (Fig. 5A), without evidence of cell damage or inflammation. Two ball pythons with fibrinonecrotic rhinitis and stomatitis (E-B1 and -B3) also exhibited a diffuse chronic tracheitis with subepithelial infiltration by lymphocytes, plasma cells, macrophages and occasional heterophils, but no further lesions in the lower respiratory tract.

The remaining snakes showed histological evidence of nidovirus-associated proliferative disease with epithelial hyperplasia in the nasal cavity, trachea, and lung, as well as mucus and inflammatory cells filling the faveolar space, consistent with our earlier findings (6). Nidovirus NP expression was seen in the pseudostratified epithelium of the primary trabeculae and in type I and type II pneumocytes of the faveolar space; the extent and distribution varied between animals.

#### Alimentary tract

In addition to the respiratory changes, several individuals (CH-A1 and -A9, CH-B6, CH-D1 and -D2, CH-C4, CH-E; carpet pythons, green tree pythons, ball pythons and a black-headed python) exhibited a severe multifocal fibrinonecrotic esophagitis (Fig. 1E, F) with abundant nidovirus-infected intact and degenerate squamous epithelial cells. The gross intestinal lesions observed in another four cases (CH-A7, and -A8, CH-B6, CH-F1) were indeed nidovirus-induced, since all exhibited a multifocal fibrinonecrotic enteritis with nidovirus NP expression in individual intact and degenerating enterocytes (Fig. 2C, D).

#### Vascular involvement, systemic spread and systemic granulomatous and/or fibrinonecrotizing disease

In all snakes with the above-described intestinal changes (CH-A7 and -A8, CH-B6, CH-F1), the mucosal lesions were found to overlie focal inflammatory processes in the intestinal wall. These were represented by multifocal (pyo)granulomatous perivascular infiltrates of macrophages and fewer heterophils and lymphocytes which were predominantly seen in the submucosa in close proximity to the gut associated lymphatic tissue (GALT) but occasionally extended into the tunica muscularis (Fig. 2C, E). These lesions contained abundant nidovirus NP in infiltrating macrophages (Fig. 2D, F). Occasional affected vessels also exhibited fibrinoid degeneration of the wall with intramural heterophil infiltration. A multifocal pyogranulomatous vasculitis and perivasculitis of small, medium-sized and large veins and arteries was also seen in the serosa of various organs, e.g. the heart, the lung and the thymus, of four snakes (CH-A7, CH-B6, CH-C3, CH-F1) (Fig. 4 C, D). Here, nidovirus NP was not only detected in macrophages of the infiltrates, but also in endothelial cells, which often appeared to be activated (Fig. 2F), and in mononuclear cells in the vessel lumina (Fig. 2F, 4A). Staining for the monocyte/macrophage marker Iba-1 showed that the mononuclear cells in the vessel lumina were predominantly Iba-1 positive, i.e. monocytes (Fig. 4B). The latter finding suggests monocyte-associated viremia, and viral spread via infected monocytes.

**Figure 4.**
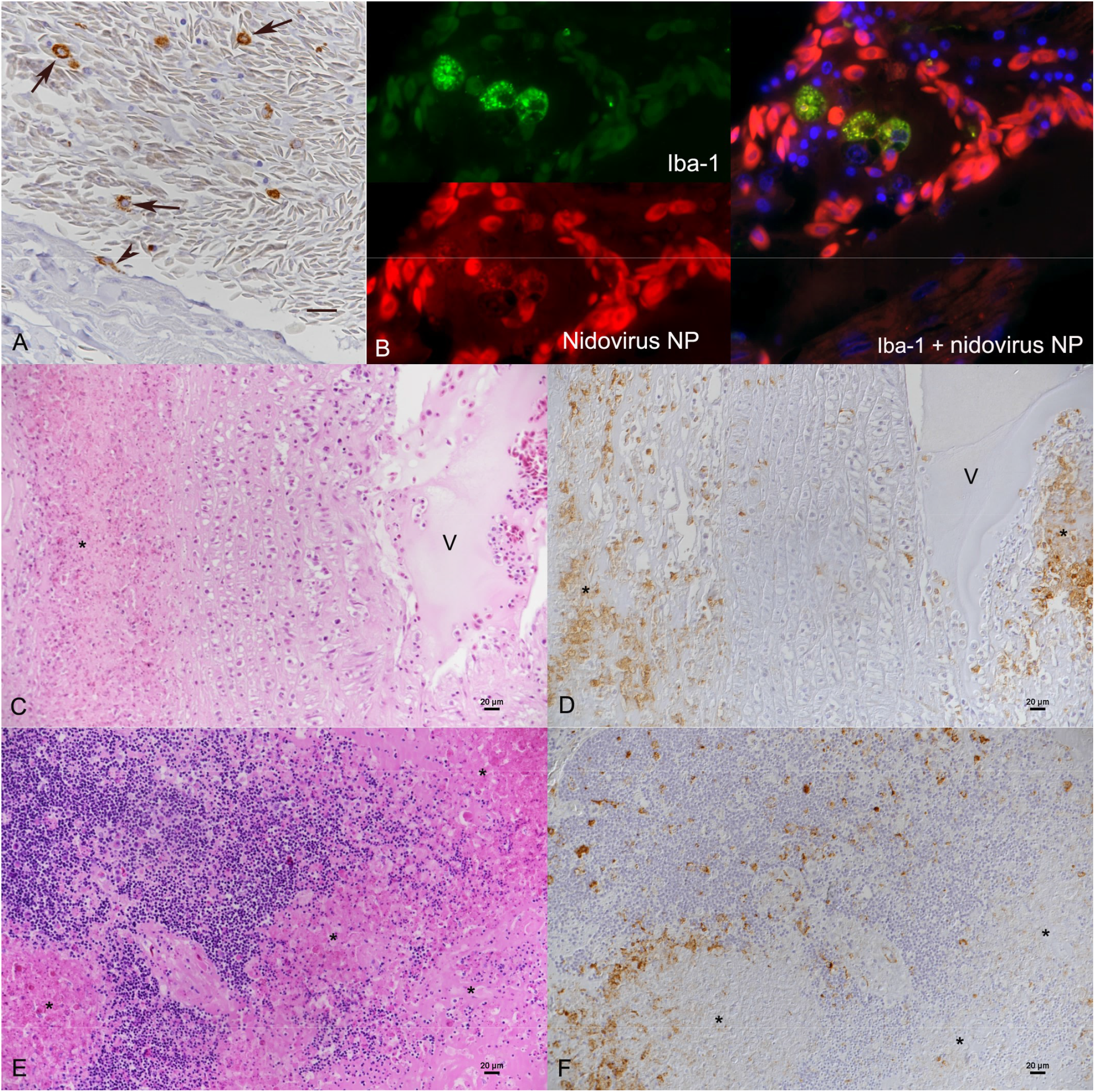
Python nidovirus disease. Involvement of blood vessels and monocyte-associated viremia. **A, B**. Snake CH-C3 (Green tree python; *M. viridis*). Lung, blood vessel. **A**. Nidovirus NP expression is seen in circulating monocytes (arrows) and individual endothelial cells (arrowhead). Immunohistology, hemalaun counterstain. **B**. Double immunofluorescence confirms nidovirus NP expression (red) in monocytes (Iba-1+; green). NB: The reddish/pinkish fluorescence seen in the abundant oval shaped cells represent autofluorescence of erythrocytes. **C, D**. Snake CH-A7 (Green tree python; *M. viridis*). Lung, serosal artery. Mild transmural mononuclear infiltration and marked granulomatous to necrotizing perivascular infiltration (asterisk; C) with abundant nidovirus NP expression (D) within infiltrating cells. V: vessel lumen. Bars = 20 μm. **E, F**. Snake CH-B6 (Green tree python; *M. viridis*). Thymus. Focal areas of necrosis (asterisks; E), surrounded by abundant nidovirus NP-positive macrophages (E). Nidovirus NP expression is also seen cell-free with in areas of necrosis. Bars = 20 μm. C, E: HE stain, D, F: Immunohistology, hemalaun counterstain.

Interestingly, all animals with the described vascular lesions were green tree pythons (CH-A7, CH-B6, CH-C3, CH-F1) and also exhibited multifocal pyogranulomatous lesions, which consisted of a central area of necrosis, surrounded by virus-laden macrophages (Fig. 4 C, D). Affected organs included the kidneys, thymus, heart and liver, and in one animal (CH-C3) the lung. One green tree python (CH-F1) also exhibited pyogranulomatous and fibrinonecrotic lesions in the oviduct, with nidovirus NP found cell-free in necrotic debris. This was seen together with a fibrinous coelomitis in the caudal coelomic cavity, adjacent to the inflamed oviduct. Pyogranulomatous lesions varied in distribution and extent between individuals, but were always present in more than one organ. There was no evidence of bacteria within the lesions. Animals with pyogranulomatous lesions often showed nidovirus NP in epithelia adjacent to the inflammatory foci (e.g. pancreatic ducts, renal tubules etc.) without any evidence of degeneration or necrosis (Fig. 5B, C).

**Figure 5.**
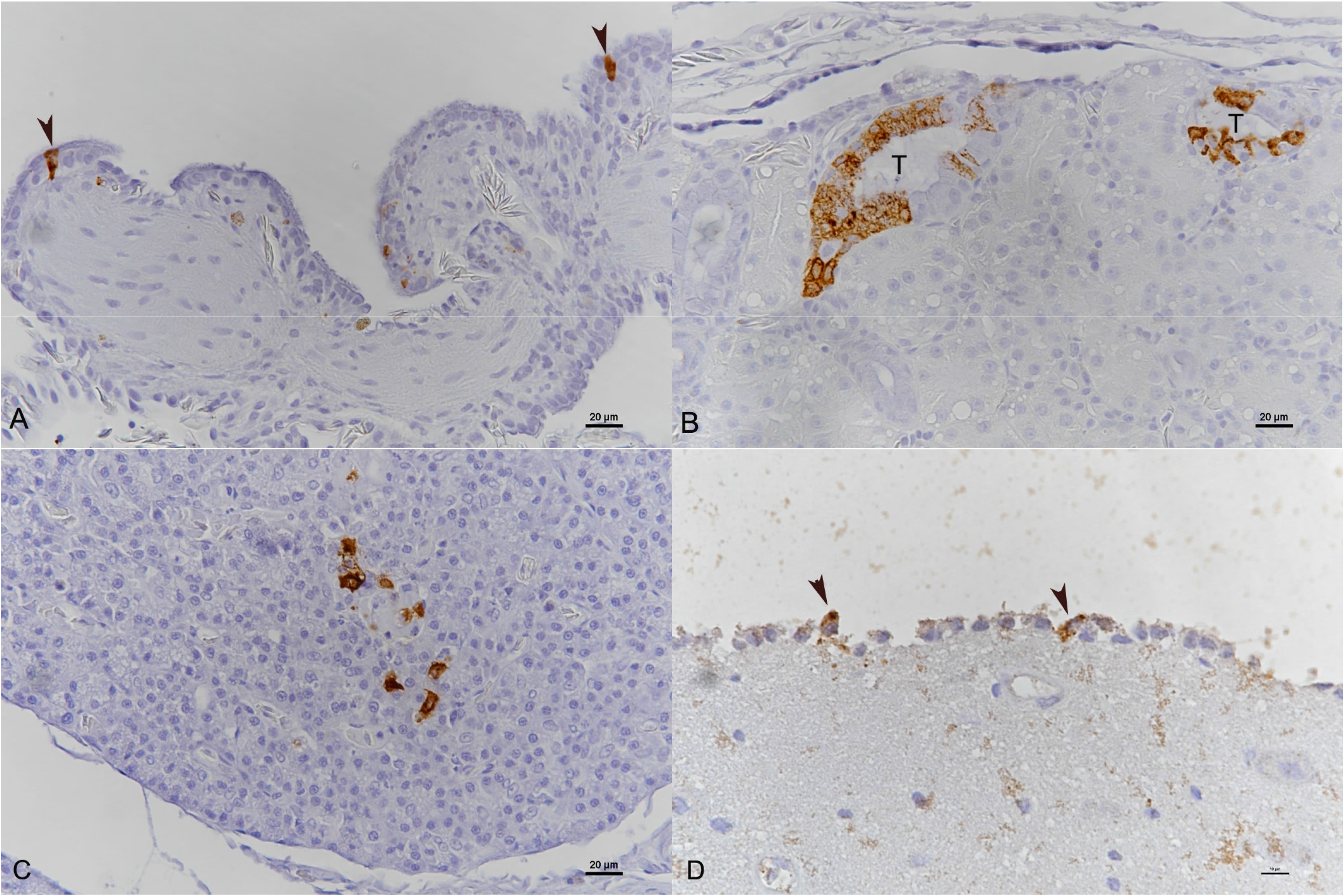
Python nidovirus disease. Nidovirus NP expression in various types of epithelial cells. **A.** Snake CH-B1. Lung (Carpet python; *Morelia spilota).* Positive respiratory epithelial cells (arrowheads). **B, C**. Snake CH-B3 (Green tree python; *Morelia viridis)*. **B.** Kidney. Positive tubular epithelial cells. T: tubular lumen. **C.** Pancreas. Nidovirus NP expression in pancreatic duct epithelial cells. **D**. Snake. CH-F1 (Green tree python; *M. viridis*). Brain. Positive ependymal cells lining the ventricle. Immunohistology, hemalaun counterstain. A-C: Bars = 20 μm, D: Bar = 10 μm.

Overall, fibrinonecrotic lesions were observed in five green tree pythons (CH-A1 and -A7, CH-B6, CH-C3, CH-F1) and one carpet python (CH-B1). They were represented by diffuse necrosis and fibrin exudation in parenchymatous organs (spleen, thymus) or on the serosal surface of the coelomic cavity (in close proximity to affected organs) with minimal inflammatory response (Fig. 4E and F). These lesions were often observed in animals with nidovirus-associated perivascular/vascular and/or pyogranulomatous lesions, indicating massive systemic spread of the virus.

#### Infection of epithelial cells and ependyma

As described, nidovirus NP was detected in various types of epithelial cells, often in close proximity to granulomatous or fibrinonecrotic lesions and without evidence of a cytopathic effect. Affected epithelial cells included type I and type II pneumocytes of the lung, epithelial cells of renal tubules and pancreatic ducts, mesothelial cells (Fig. 5A-C) and hepatocytes (Fig. 3D). In two cases (CH-B6 and –F1) ependymal cells were also found to be infected, without evidence of correlating brain lesions (Fig 5D).

### Virus isolation and identification

Nidovirus isolation was attempted by inoculating primary cell cultures of green tree python fetal liver and brain tissue with lung tissue homogenates of 24 animals that had been confirmed as nidovirus infected by immunohistology and RT-PCR. After inoculation, cytopathic effects were observed in 12/24 samples at about 3-4 days post infection (dpi). Two nidovirus positive lung (CH-A8, CH-F2, Table 1) and three positive liver specimens (CH-A7, –A8 and CH-B6, Table 1), a lung lavage sample, and 16 cell culture supernatants (Table 1) were subjected to NGS. To investigate if the viruses were of the same species, we included samples from both Swiss and Spanish collections. The approach yielded nidovirus sequences from 14 lung culture supernatants. *De novo* genome assembly from tissue sample after removal of reads matching to the host’s genome produced contigs covering the complete coding sequence (CDS) of a nidovirus novel in Switzerland. The genome organization of the identified nidovirus was identical to those described for Ball python nidovirus (BPNV) and Morelia viridis nidovirus (MVNV). The open reading frame 1b (ORF1b, RNA-dependent RNA polymerase) of these viruses had less than 5 % nucleotide and less than 2.4 % amino acid differences with recently described Morelia viridis nidovirus isolates BH128 14-12 and BH171 14-7 and Serpentovirinae sp isolates L3, L4 and H0-1, but more than 13 % nucleotide and 8.5 % amino acid differences to all other nidovirus strains. Phylogenetic analysis based on ORF1b suggested that these virus forms a sister clade to the previously known python-associated nidovirus species; BPNV and MVNV (Fig 6). CDSs of the same virus species were obtained from 11 cell culture supernatants, including the isolation attempts from the liver samples included in the NGS analysis. One of the cell culture supernatants analyzed by NGS yielded CDS of MVNV (animal E-C1, Table 1), and for four cell culture supernatants NGS failed due to severe bacterial contamination.

**Figure 6.**
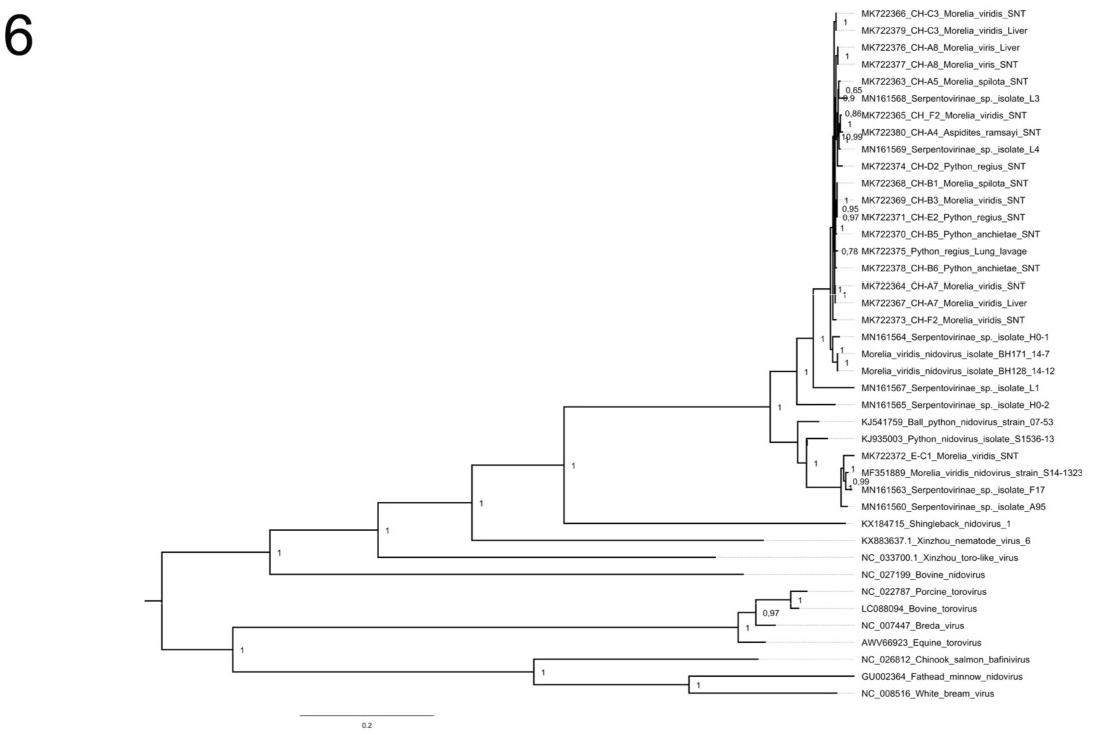
Maximum clade credibility tree constructed from the ORF1b amino acid sequences of representatives of the family Tobaniviridae. The phylogenetic tree was constructed using the Bayesian MCMC method with the WAG model of substitution. Posterior probabilities are shown at each node.

For three snakes (CH-A7-8, CH-C3, Table 1) the analysis included cell culture isolated virus (lung homogenate as inoculum) and virus sequenced directly from tissue (liver). The consensus sequences obtained from liver tissue and the respective cell culture isolate were identical for each of these three snakes indicating that isolation did not induce a marked bias in the sequence analysis. Single nucleotide variant (SNV) analysis of the NGS data did not show evidence of significant differences in the sequences obtained from different organs or from cell culture isolates. Likewise, the SNV analysis did not either find differences between the viral populations obtained from animals with or without systemic infection, suggesting that viral intra-host polymorphism does not explain the varying pathogenic manifestation.

### Recombination

The initial NeighborNet analysis of snake-associated nidoviruses suggested strong evidence of recombination (PHI test p<0.001) (Fig 7A). Similarly, the tree order scan suggested a high amount of phylogeny violations across the genome. Therefore, we conducted a more detailed recombination analysis. This analysis suggested at least five highly supported recombination events.

**Figure 7.**
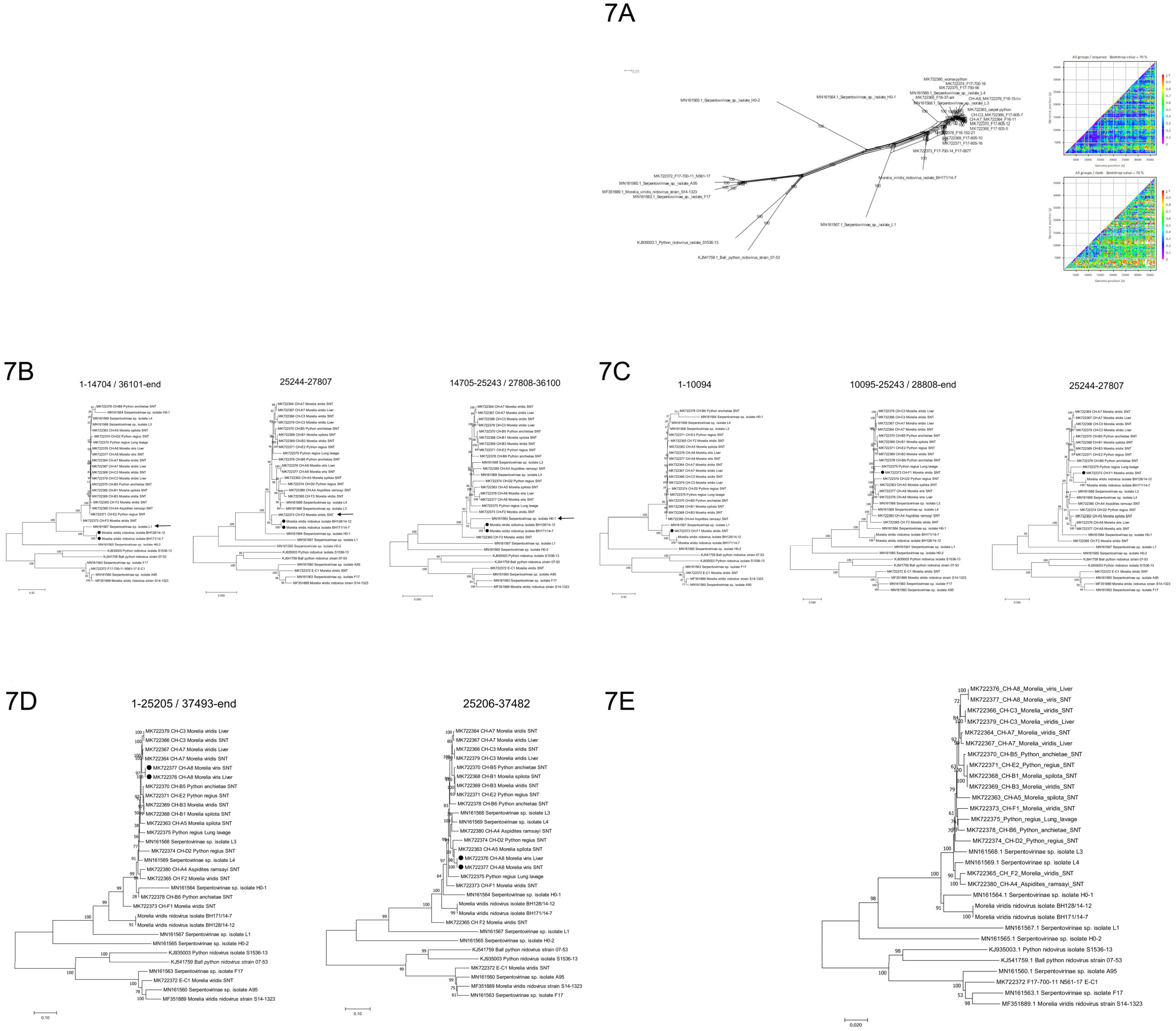
Recombination analysis of nidovirus genomes. (A) Phylogenetic network and tree order scan showed evidence of recombination in the sequence data set. (B-D) Incongruent tree topologies of virus strains with highly supported recombination events estimated using RDP software. The numbers above trees show the genome regions in alignment. (E) Phylogenetic tree based on concatenated non-recombinant genome regions.

Morelia viridis nidovirus isolates BH171/14-7 and BH128/14-12 (33) cluster together with Serpentovirinae sp. isolate L1 based on 5’ and 3’ ends of the genome (corresponding to nucleotides 1-14704 and 36100-37869 in the alignment), whereas based on nucleotides 14705-25143 and 27808-36099 these strains cluster together with Serpentovirinae sp. isolate H0-1 (MN161564), and between nucleotides 25144 and 27807 together with strain CH-F1 (MK722373) (Fig 7B). Likewise, strain CH-F1 clusters together with Serpentovirinae sp. isolate L1 and BH171/14-7 on the basis of nucleotides 1-10094 but together with MK722375_Python_regius based on the 3’ end of the genome (Fig 7C). This suggests that the Serpentovirinae sp. isolate L1 (MN161567) has been involved in at least two distinct recombination events. Although these four isolates (Serpentovirinae sp. isolate L1, Morelia viridis nidovirus isolates BH171/14-7 and BH128/14-12 and strain CH-F1) cluster together in the 5’end of the genome, the genetic distances between these strains are rather high in this region, suggesting that the exact recombination partners are not represented in the dataset. Since Serpentovirinae sp. isolate L1 was sequenced from a *Python regius* in the USA in 2015 (14), whereas the strains BH171/14-7, BH128/14-12 (33) and CH-F1 were isolated from a *Morelia viridis* in Germany (2015) and Switzerland (2018), it is likely that the recombination events have occurred between the ancestors of these lineages.

Another clear recombination event was detected in the strain CH-A8 (MK722376), which clusters together with the strains CH-A7 and CH-C3 based on the majority of the genome, whereas between nucleotides 25205-37482 it clusters with CH-A5 (MK722363) (Fig 7D). In addition, the strain CH-D2 clusters together with this group (CH-A8 / CH-A5) based on nucleotides 25205-37482.

Sequential exclusion of recombinant regions from the alignment (i.e. nucleotides 1-14704 and 25144-37869) resulted in the tree shown in Figure 7E. Although some of the methods included in RDP4 analysis still found evidence of recombination in this alignment, the further removal of these regions did not affect the topology of the tree.

The phylogenetic tree indicated that multiple viral strains circulate among the snakes of a single breeder and, on the other hand, there are some indications of viral transmission between the breeders, as exemplified by the high similarity of strains CH-B1, CH-B3 (breeder CH-B) and CH-E2 (breeder CH-E).

The breeders CH-A, CH-B, CH-C and CH-F exchanged animals, which may explain the grouping of strains CH-C3, CH-A7, CH-E2, CH-B1, CH-B3, CH-B5 and CH-A8 to a single cluster. Notably, no recombination events were detected between the newly detected nidovirus species and the previously characterized Morelia viridis nidovirus, python nidovirus and ball python nidovirus (Fig 7E), suggesting that recombination occurs only within virus species but not between species.

### Evidence of horizontal transmission within colonies and of virus shedding from diseased animals

One Swiss breeder (CH-B, Table 1) lost one of three juvenile carpet pythons from the same clutch that had been housed separate from each other, but in the same room (distance less than 0.5 m), to severe fatal proliferative pneumonia (CH-B4). The post mortem examination confirmed nidovirus-associated pneumonia in the deceased animal, prompting the breeder to submit the two siblings (CH-B1 and -B2, Table 1) for euthanasia and diagnostic post mortem examination. Both individuals exhibited nidovirus-associated lesions in the upper airways but not in the lung, suggesting that these cases represented an early disease stage following airborne infection.

In addition to the above snakes, we studied choanal and/or cloacal swabs from seven additional animals from different collections for the presence of nidovirus RNA by RT-PCR. All of the tested animals were found positive for nidovirus infection (Table 1), and included individuals with nidovirus-associated proliferative disease (respiratory form) and individuals with evidence of granulomatous and fibrinonecrotic disease (systemic form), suggesting that diseased animals shed the virus. Further, the virus strains from snakes CH-B1, CH-B3 and CH-B5 showed high genetic identities and clustered together in the phylogenetic trees suggesting viral transmission between the snakes of breeder CH-B.

### Pathological changes unrelated to nidovirus infection

Individual snakes exhibited additional lesions that appeared to be unrelated to nidovirus infection, as confirmed by immunohistology for nidovirus NP; renal gout (CH-A3 and E-A1, Table 1), and a necrotzing splenitis (CH-A1 and CH-C4, Table 1) and fibrinonecrotic enteritis with intralesional coccoid bacterial colonies (E-B1, Table 1). Both conditions are likely a consequence of debilation/dehydration due to chronic pulmonary disease.

## DISCUSSION

Nidoviruses have recently been described as the cause of respiratory tract disease in several python species in the USA and Europe (5, 6, 10, 11), representing an emerging threat to python traders and breeders in particular. The present study aimed to further elucidate the characteristics and relevance of python nidoviruses. Particular emphasis was laid on the potential species specificity of and/or susceptibility to the viruses, their phylogeny and geographic divergence as well as viral shedding and transmission, host cell tropism and type of associated disease. Experimental infection has been shown to reproduce the naturally occurring respiratory disease (7), so our study focused on natural cases from different geographic regions, from breeding colonies that often kept several python species, and from colonies that at least partly worked closely together, exchanging animals.

Pythons are nonvenomous snakes found in Africa, Asia and Australia. Eight genera and 31 species are currently recognized, most are also kept in captivity (34). The present study shows that python species originating from three continents are susceptible to python nidoviruses and can develop nidovirus-associated disease. Ball pythons (*Python regius* (35)) and Angolan pythons *(Python anchietae* (36)) are closely related and originate from sub-Saharan and southern Africa, respectively. The green tree python (*Morelia viridis* (Schlegel, 1872)), the carpet python (*Morelia spilota* (37)) and the Ramsay’s python (*Aspidites ramsayi*, (38)) represent species originating from Australia and New Guinea (Indonesia and Papua New Guinea), whereas the Indian python (*Python molurus* (39)) is indigenous to the Indian Subcontinent and Southeast Asia.

In our study, no differences were noted between the python species in the degree and distribution of nidovirus-associated respiratory disease. However, it is interesting to note that all animals with systemic viral spread (indicated by disseminated granulomatous and/or fibrinonecrotic lesions, [peri]vascular lesions, infected monocytes etc.) were of the genus *Morelia*. The results suggest increased susceptibility of snakes from this genus to nidovirus infection and/or disease, a finding also supported by recent epidemiologic studies (14). In support, the analysis of the viral genomes by NGS did not show differences in the viral genome that would explain the differences in the disease manifestation. One could speculate that each python species has its own nidovirus, and that cross-species transmission of the viruses results in morbidity. Should pythons (or snakes in general) indeed be hosts for nidoviruses, then a co-infection with an unidentified bacterial or viral agent could also explain the sudden emergence of fatal pneumonia cases in captive snakes. An alternative explanation for the emergence of nidovirus infections in python collections could be that there have been multiple nidovirus introductions to various python species from an unknown host species or a vector.

Recently, experimental infection of ball pythons established the causal relationship between nidovirus infection and mucinous inflammation of the upper respiratory and the gastrointestinal tract as the main pathological process early, i.e. within the first 12 weeks, after infection (7). Infected pythons showed severe respiratory distress with only minimal pneumonia, most likely because the mucus overproduction resulted in obstruction of the upper airways (7). These findings correlate with those we made when studying naturally infected cases; some animals were submitted for euthanasia due to acute respiratory distress, but only exhibited inflammatory processes in nasal and oral cavity, without histologic evidence of tracheitis or pneumonia. In all other naturally infected animals, rhinitis, stomatitis and tracheitis were present, as well as a variable degree of proliferative pneumonia, indicating longer duration and a more progressed stage of the disease. On that note, the anatomic structure of the snake lung has to be considered, as studies on Burmese pythons have shown that python lungs provide excess capacity for oxygen exchange. This leads to progressive spread of respiratory infections through the lung, thereby continuously reducing respiratory gas exchange without causing clinical signs. The latter will develop only when the oxygen exchange capacity falls below the requirements of the metabolic rate. This is particularly relevant in association with hyperplasia of the pulmonary epithelium, which has an impact on the blood gas exchange (40). In some snakes, we additionally observed an esophagitis; this is interpreted as a consequence of overspill and swallowing of virus-laden mucus, a theory also supported by previous studies (6, 7, 40). It is possible that the esophagus is involved because its epithelium contains ciliated cells in various snake species (13), and ciliated cells represent the primary site of viral replication, a feature also shown for human coronaviruses (13, 41, 42).

Similar to python nidoviruses, mammalian toroviruses (found in horses, swine and cattle) belong to the family *Tobaniviridae* and have a worldwide distribution (43). Epidemiologic studies are sparse, yet seroprevalence seems to be high in affected populations. For example, the seroprevalence to porcine torovirus (PToV) exceeds 95 % in swine populations, and is 94 % in cattle (Breda virus) (44) and 38% in horses (Berne virus) (45). Neutralizing antibodies to Berne virus (equine torovirus) have also been detected in goats, pigs, rabbits, and some species of wild mice. Equine and porcine toroviruses are generally associated with asymptomatic enteric infections, and transmission is probably via the oral/nasal route through contact with feces (46) or nasopharyngeal secretions (47). However, torovirus infection has been associated with neonatal and postweaning diarrhea in swine (48) and severe diarrheal disease in cattle (49). In our study nidovirus RNA was detected in choanal and cloacal swabs of individuals presenting with either the respiratory (local) or systemic form of the disease. This correlates with studies on human SARS-CoV-2, where viral RNA was detected not only in throat swabs but also in stool samples (50). Viral shedding via the feces in pythons without intestinal lesions could likely be a consequence of the swallowing of mucus. Therefore, besides the respiratory (aerosol) route, the fecal-oral route also appears to be a likely way of transmission in pythons. The literature suggests that mammalian coronaviruses have a limited host cell tropism and affect either the intestinal or the alveolar epithelium (6, 7, 10, 11, 47, 51–53). This seems not to apply to python nidoviruses for which the results of the present study indicate a rather broad cell tropism, for various types of epithelia (respiratory and pulmonary epithelium, enterocytes, hepatocytes, epithelial cells in renal tubules and pancreatic ducts) and ependymal cells. Interestingly, a similar tendency is described for SARS-CoV-2 (54). Previous studies on python nidoviruses have hinted at a broader target cell spectrum, but did so far not go beyond the detection of nidovirus RNA in organs other than the respiratory tract (10), and did neither identify infected cells nor correlate infection with actual organ lesions.

Of particular interest is the fact that python nidoviruses also infect non-epithelial cells distributed over the entire body. We detected nidovirus NP in endothelial cells, intravascular monocytes and extravascular macrophages within inflammatory processes. The latter suggest that monocytes facilitate the systemic spread of the virus. So far, viruses of the subfamily *Tobaniviridae* have not been reported to infect monocytes/macrophages. However, monocyte/macrophage infection is known to be a key process in the pathogenesis of other members of the order *Nidovirales*, such as feline coronaviruses (FCoV) (55), ferret systemic coronavirus (FRSCV) (56) and Severe Acute Respiratory Syndrome Coronavirus (SARS-CoV) (57). For FCoV, the ability to infect, replicate in and activate monocytes and macrophages is essential in the pathogenesis of feline infectious peritonitis (FIP), a fatal disease of felids. While the low-virulence feline enteric coronavirus (FECV) biotype primarily replicates in enterocytes and does only induce mild enteric disease, the highly virulent feline infectious peritonitis virus (FIPV) biotype predominantly arises after S gene mutations in FECV of the infected host that allow efficient replication in and systemic spread with monocytes (58). This allows rapid dissemination of the virus throughout the body (monocyte-associated viremia) and is a prerequisite of the monocyte activation with subsequent development of the granulomatous phlebitis that is the hallmark of FIP (55, 59, 60). A similar mechanism is likely for ferrets with a comparable disease and for experimental infections of IFN-γ knock-out mice with Mouse Hepatitis Virus (MHV) (56, 61).

Similar to cats with FIP, systemically nidovirus-infected pythons exhibited a multifocal macrophage-dominated vasculitis which was in severe cases associated with fibrinoid necrosis of the vessel wall, but also appeared as chronic perivascular (pyo)granulomatous cuffs with abundant nidovirus-positive macrophages. These vascular lesions could result from an interaction between activated nidovirus-infected monocytes and endothelial cells, in particular since the latter were also found to become infected. They might also be responsible for the broad spectrum of granulomatous to fibrinonecrotic lesions in organs, since these might at least partly be of ischemic nature.

Sequencing of the nidovirus genomes did not reveal variants that would be associated with the systemic infection observed in some individuals. The genome length and relatively high mutation and recombination rate of nidoviruses makes identification of mutations that might alter the pathogenesis of the virus rather challenging. The S protein is the precursor of the spike complex that mediates receptor binding and entry, and thus mutations in the S protein could most easily explain the differences in tissue tropism (62). However, we did not identify S protein mutations that would explain the different phenotypes. One could thus speculate that the different infection outcome is a consequence of differences in the host immune response.

During viral infections of mammals, viruses are recognized by pattern recognition receptors (PRRs) which induce a type I interferon (IFN) response, mediated by IFN-α and -β. The IFN response is considered the primary host defense mechanism against viral infections. It is therefore not surprising that many viruses have developed mechanisms to subvert or alter the type I IFN response (61). Interference of nidoviruses with the IFN response has so far mainly been investigated in the family *Coronaviridae*. In studies on human SARS-CoV infection the presence of viral proteins in monocytes was not found to be associated with significant IFN-α production and it was suggested that viral particles had been taken up by monocytes via phagocytosis. This correlates with studies on IFN receptor deficient knock-out mice where MHV infection was associated with higher viral titers and a broader tissue tropism of the virus (61). Further studies are needed to determine whether systemic nidoviral infection in snakes is also related to an altered interferon response.

## ACKNOWLEDGEMENTS

The authors are grateful to the technical staff of the Histology Laboratory for excellent technical assistance. The study was supported by the Academy of Finland (1308613) and the Finnish Foundation of Veterinary Research. We also wish to thank the snake owners for submitting the cases.

